# Opposite regulation of glycogen metabolism by cAMP produced in the cytosol and at the plasma membrane

**DOI:** 10.1101/2022.09.01.505928

**Authors:** Arthur J. Verhoeven, Paulo F.V. Bizerra, Eduardo H. Gilglioni, Simei Go, Hang Lam Li, Ronald P.J. Oude Elferink, Jung-Chin Chang

## Abstract

Cyclic AMP is produced in cells by two different types of adenylyl cyclases: at the plasma membrane by the transmembrane adenylyl cyclases (tmACs, *ADCY1∼ADCY9*) and in the cytosol by the evolutionarily more conserved soluble adenylyl cyclase (sAC, *ADCY10*). By employing high-resolution extracellular flux analysis to study glycogen breakdown in real time, we show here thatcAMP regulates glycogen metabolism in opposite directions depending on its location of synthesis within cells. While the canonical tmAC-cAMP-PKA axis promotes glycogenolysis, we demonstrate that the non-canonical sAC-cAMP-Epac1 signalling suppresses glycogenolysis in a variety of cell types. Our findings demonstrate the importance of cAMP microdomain organization in glycogen metabolism and reveal a novel role of sAC in energy metabolism during glucose deprivation.

## Introduction

Since the discovery of 3’,5’-cyclic adenosine monophosphate (cAMP) as the second messenger in hormone-mediated glycogenolysis (Berthet et al., 1957; Sutherland and Rall, 1957; Sutherland and Rall, 1958), nine transmembrane adenylyl cyclases (tmACs, *ADCY1*∼*ADCY9*) has been identified as sources of this versatile second messenger at the plasma membrane (reviewed in (Hanoune and Defer, 2001)). Much later, a distinct type of mammalian adenylyl cyclase, soluble adenylyl cyclase (sAC, *ADCY10*), was discovered in the cytosol (Buck et al., 1999). sAC is evolutionarily more conserved than tmACs and is not localized to the plasma membrane but exclusively to the cytoplasm (Chen et al., 2000; Kamenetsky et al., 2006). cAMP produced in the cytosol by sAC and at the plasma membrane by tmACs share the same cAMP effectors, including protein kinase A (PKA) and “exchange factor directly activated by cAMP” (Epac1 and Epac2). However, in contrast to tmACs, sAC is neither regulated by G-proteins nor activated by forskolin, a broad-spectrum tmAC activator (Buck et al., 1999; Chen et al., 2000). Rather, sAC is activated by bicarbonate and ATP in the physiological ranges and fine-tuned by the free Ca^2+^ concentration (Chen et al., 2000; Geng et al., 2005; Kleinboelting et al., 2014; Litvin et al., 2003; Zippin et al., 2013).

While the tmACs are evolved in multicellular organisms and are activated or inhibited by signalling of hormones and neurotransmitters to mediate intercellular communication, the bicarbonate-activable sAC homologs are conserved in both multicellular and unicellular organisms and serve more cell-autonomous functions (Fredriksson and Schioth, 2005; Kamenetsky et al., 2006; Kobayashi et al., 2004). Firstly, sAC regulates oxidative phosphorylation by sensing bicarbonate derived from CO_2_ produced by the tricarboxylic acid cycle (Acin-Perez et al., 2009) and free Ca^2+^ in the mitochondrial matrix (Di Benedetto et al., 2013). Secondly, sAC is an ATP sensor due to its high *K*_m_ for ATP (ranging from 1 to 10 mM) (Jaiswal and Conti, 2003; Litvin et al., 2003; Zippin et al., 2013). Thirdly, Ca^2+^, another versatile and universal second messenger, stimulates sAC in synergy with bicarbonate (Geng et al., 2005; Jaiswal and Conti, 2003), allowing sAC to fine-tune cellular metabolism. Importantly, sAC is expressed in almost all tissues examined (Geng et al., 2005; Levin and Buck, 2015). We have recently shown that sAC acts as an acute switch for aerobic glycolysis and oxidative phosphorylation maintaining the autonomous energy metabolism of cells (Chang et al., 2021).

Over the past two decades, accumulating evidence supports that the versatility of cAMP signalling is based on the compartmentalization of cAMP signalling (Agarwal et al., 2022; Kamenetsky et al., 2006; Lefkimmiatis and Zaccolo, 2014). From its site of generation, cAMP concentration decays and forms a signalling microdomain due to buffering by regulatory units of PKA (Agarwal et al., 2016) and degradation by PDEs (Jurevicius and Fischmeister, 1996; Zaccolo and Pozzan, 2002). The spatial specificity is further enhanced by scaffolding proteins, such as A-kinase anchoring proteins (Pidoux and Tasken, 2010), that assemble adenylyl cyclases, PDEs, and cAMP effectors at the same location. Indeed, cAMP generated at the plasma membrane by tmACs and cAMP generated in the cytosol by sAC or bacterial adenylyl cyclases can exert opposite regulation of apoptosis (Chang et al., 2016; Kumar et al., 2009) and endothelial barrier function (Obiako et al., 2013; Sayner et al., 2004). Although it is well-established that tmAC-derived cAMP activates PKA, which induces a phosphorylation cascade of phosphorylase kinase and glycogen phosphorylase to induce glycogenolysis (Berthet et al., 1957; Fischer and Krebs, 1955; Sutherland and Rall, 1957; Walsh et al., 1968), sAC-derived cAMP has also been implicated in affecting glycogen levels in astrocytes (Choi et al., 2012; Jakobsen et al., 2021). However, it is unknown whether sAC-derived cAMP and tmAC-derived cAMP regulate glycogen metabolism by different mechanisms and, if so, how the specificity of cAMP signalling is maintained in the same cell. In the present study, we applied high-resolution extracellular flux analysis to study glycogen breakdown in real-time and revealed opposite regulation of glycogen metabolism by sAC-derived cAMP and tmAC-derived cAMP. While the well-established tmAC-cAMP-PKA signalling axis promotes glycogenolysis, the hitherto unknown sAC-cAMP-Epac1 signalling axis inhibits glycogenolysis.

## Results and Discussion

### Inhibition of soluble adenylyl cyclase in the absence of glucose causes a transient stimulation of glycolysis in HepG2 cells

To investigate the possible role of sAC in glycogen metabolism, we first made use of the Seahorse flux analyzer which measures in real time not only the oxygen consumption rate (OCR) of cells, but also the extracellular acidification rate (ECAR). Although both glycolysis and TCA cycle activity contribute to ECAR, glycolytic flux accounts for the majority of ECAR (Mookerjee et al., 2017; Mookerjee et al., 2015). ECAR thus represents a good first estimation of glycolytic flux. In the absence of added glucose, glucose units for glycolysis may be generated by glycogen breakdown. We therefore reasoned that ECAR might be a good indicator of glycogenolysis in the absence of added glucose. Indeed, when HepG2 cells were incubated with octanoate (a membrane-permeable substrate for mitochondrial β-oxidation) and subsequently stimulated with the tmAC activator forskolin, we observed a clear increase in ECAR that was acute and transient in nature (Figure 1A), which is consistent with acute increase of glycolytic flux by transient fuelling of glycogen-derived glucose units until the glycogen store was depleted. Interestingly, the transient change in ECAR was accompanied by a reciprocal, transient decrease in OCR (Figure 1B). Surprisingly, when acutely inhibiting sAC-derived cAMP with the sAC-specific inhibitor LRE1 (Ramos-Espiritu et al., 2016), a transient increase in ECAR with a reciprocal decrease in OCR was also induced (Figure 1C and 1D). This indicated that acute decrease of sAC-derived cAMP caused a transient increase in glycolytic flux during glucose deprivation. Indeed, when HepG2 cells were incubated in the presence of 2-deoxyglucose (2-DG, an inhibitor of glycolysis), the transients induced by sAC inhibition were abolished (Figure 1E and 1F). Calculation of the concomitant ATP production rates showed that ATP production by glycolysis was transiently stimulated upon sAC inhibition, while ATP production by TCA cycle and oxidative phosphorylation were transiently suppressed (Figure S1A-S1D). During these transient changes in ECAR and OCAR, the overall ATP production (Figure S1E) and the adenylate energy charge (Figure S1F) remained constant, indicating that sAC co-ordinates ATP production of glycolysis and oxidative phosphorylation to maintain energy homeostasis not only during glucose-sufficient conditions (Chang et al., 2021), but also during glucose deprivation. Since glycogen is the primary glucose reserve in cells during glycose deprivation, we tested the hypothesis that in the absence of glucose sAC inhibition mobilized glycogen to cause a transient increase in glycolytic flux. As a first approach to investigate this new mechanism of glycogen regulation by cAMP, we tested whether CP-91149, an established inhibitor for glycogen phosphorylase *a* (*i*.*e*. with a phosphorylated Ser-15 residue) (Martin et al., 1998), could prevent the transient ECAR increase upon acute sAC inhibition. Indeed, we observed that the changes in ECAR and OCR induced by sAC inhibition were largely prevented by co-incubation with CP-91149 (Figure 1G and 1H). These results suggested that sAC-dependent cAMP signalling suppresses glycogen metabolism, which is opposite to the established glycogenolytic effect of tmAC-dependent cAMP signalling.

**Figure 1.**
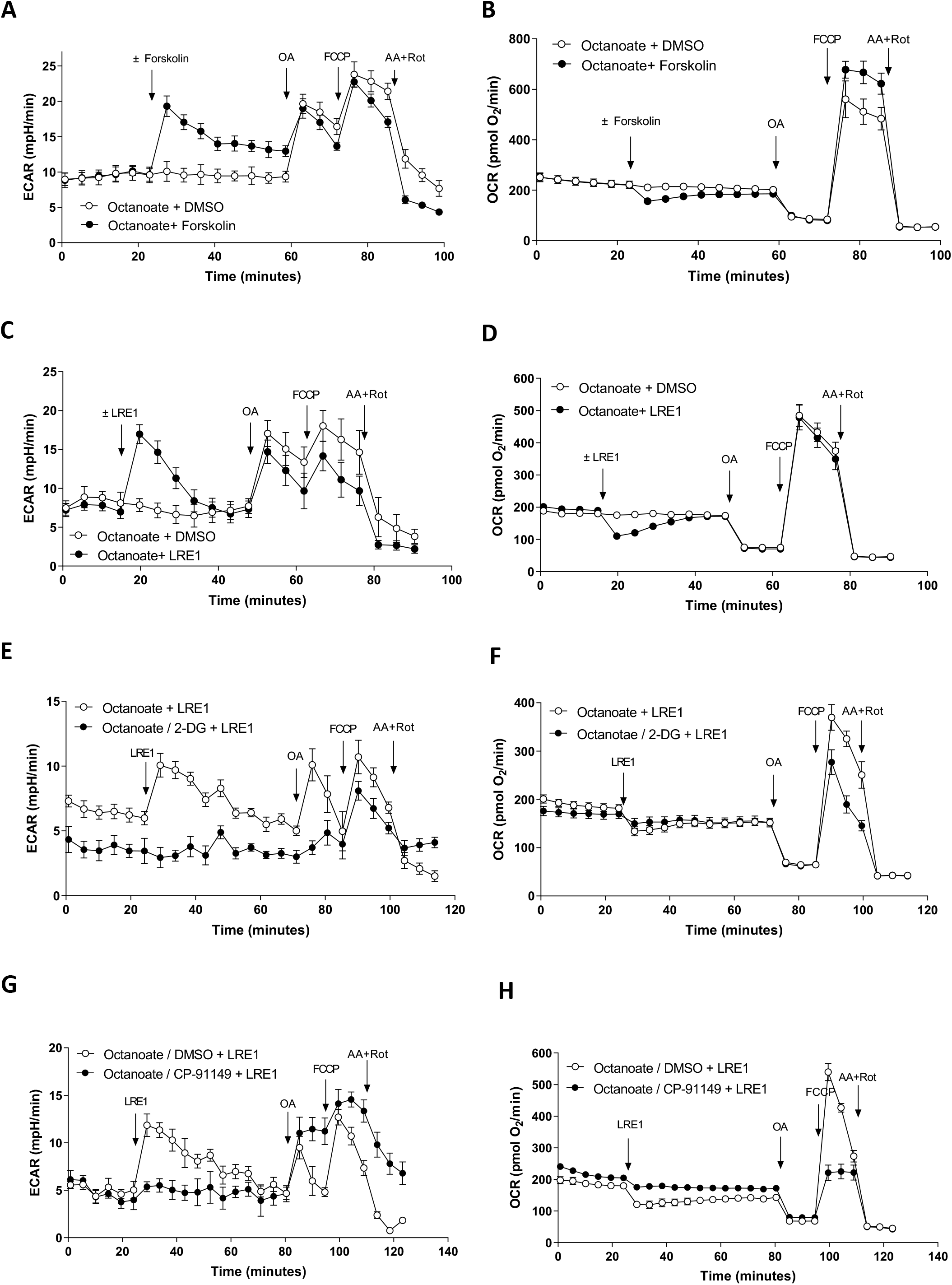
Inhibition of soluble adenylyl cyclase causes transient changes in glycolysis and oxygen uptake in HepG2 cells. HepG2 cells were preincubated for 1 hour in HBSS for ambient air in the presence of 125 µM octanoate. The extracellular acidification rate (ECAR) and the oxygen consumption rate (OCR) were then analyzed by Seahorse Flux Analyzer XF96. Arrows indicate the injection of 0.1% DMSO (vehicle control) or 1 µM forskolin (A-B) or 50 µM LRE1 (C-H). Subsequently, oligomycin A (OA), carbonyl cyanide-*p*-trifluoromethoxy-phenyl hydrazone (FCCP), and antimycin A (AA) and rotenone (Rot) were added as indicated in the figures. HepG2 cells were also tested in the presence of 5,5 mM 2-deoxyglucose (2-DG, E-F) or 100 µM CP-91149 (G-H). Data are presented as mean ± SD of at least 6 replicates for each condition. All data are representative of at least 3 independent experiments.

### Inhibition of soluble adenylyl cyclase promotes glycogenolysis in various cell types

While liver and muscles are the primary organs for glycogen storage, glycogen can be detected in virtually all cells including tumour cell lines of various tissue origins (Rousset et al., 1981). Indeed, we found that HepG2 cells store significant quantities of glycogen which were, in the absence of added glucose, greatly reduced upon sAC inhibition and upon stimulation of tmACs with forskolin (Figure 2A). The marked disappearance of glycogen under both conditions was, as expected, accompanied by a stimulation of lactate release (Figure S2). The strong decrease in glycogen content in HepG2 cells induced by sAC inhibition was not observed in the presence of glucose (Figure 2B) and was strongly diminished in the presence of the phosphorylase *a* inhibitor CP-91149 (Figure 2C), indicating that phosphorylase was activated by sAC inhibition. In subsequent experiments we investigated whether the glycogenolysis induced by sAC inhibition occurred also in other cell types. In primary mouse hepatocytes, sAC inhibition caused glycogen breakdown both in the presence of added glucose (Figure 2D) and in the absence of glucose (Figure 2E). In primary mouse hepatocytes, the glycogenolytic effect of sAC inhibition was, just as in HepG2 cells, sensitive for phosphorylase inhibition (Figure 2E). To strengthen our findings in HepG2, we examined if sAC also regulates glycogen metabolism in immortalized normal human intrahepatic cholangiocytes H69 (hereafter H69 cholangiocytes), which have a high activity of sAC (Chang et al., 2016). In H69 cholangiocytes, sAC inhibition caused net glycogen breakdown both in the presence and absence of extracellular glucose (Figure 2F). Importantly, in H69 cholangiocytes, stimulation of tmACs by forskolin and inhibition of sAC also both induced glycogen breakdown, demonstrating again the opposite effects of cAMP signalling by tmACs-derived cAMP and sAC-derived cAMP within the same cell (Figure 2G). Taken together, these data indicate sAC activity prevents rapid depletion of glycogen in the absence of glucose, which is opposite to that of the well-established tmAC-cAMP-PKA axis. Moreover, the induction of glycogenolysis upon sAC inhibition in H69 human cholangiocytes and primary mouse hepatocytes in the presence of glucose demonstrates that, at least in certain cell types, sAC-regulated glycogen turnover plays a role in determining the steady state glycogen level.

**Figure 2.**
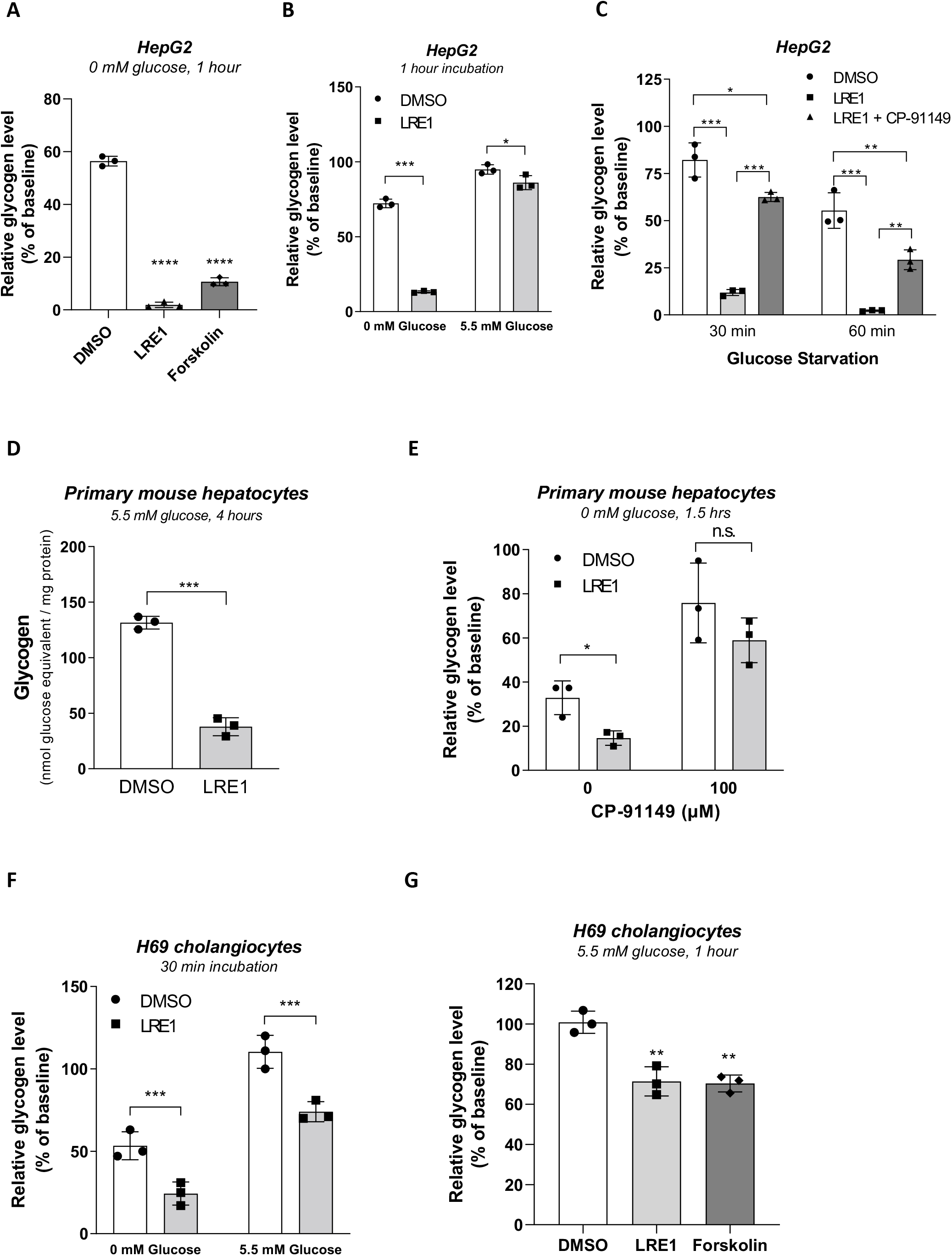
Inhibition of soluble adenylyl cyclase promotes glycogenolysis. (A) HepG2 cells were preincubated with 5.5 mM glucose for 1 hour. After harvesting baseline samples, cells were incubated with medium without glucose in the presence of 0,1% DMSO (vehicle control), 50 µLRE1 or 1 µM forskolin for another hour and then harvested for glycogen determination. Data were expressed as % of the baseline value (154 ± 21 nmol glucosyl unit per mg protein, n = 6) and mean ± SD. (B) HepG2 cells were preincubated with 5.5 mM glucose for 1 hour. After harvesting baseline samples, cells were refreshed with medium ± 5.5 mM glucose ± 50 µM LRE1 for another hour and then harvested for glycogen determination. Data were expressed as % of the baseline value (231 ± 22 nmol glucosyl unit per mg protein, n = 3) and mean ± SD. (C) HepG2 cells were preincubated with 5.5 mM glucose for 1 hour. After taking baseline samples, cells were exposed to medium without glucose in the presence of 0.1% DMSO, 50 µM LRE1, or 50 µM LRE1 + 100 µM CP-91149. Glycogen contents were determined after 30 and 60 min of incubation. Data are expressed as % of the baseline value (119 ± 2 nmol glucosyl unit per mg protein, n=4) and mean ± SD. (D) Primary mouse hepatocytes were incubated in the presence of 5.5 mM glucose for 4 hours with 50 µM LRE1 or 0.1% DMSO (vehicle control) present. Glycogen contents were determined and normalized to protein content. Data represent mean ± SD of triplicate determinations. (E) Primary mouse hepatocytes were acutely starved in the presence or absence of 50 µM LRE1 and 100 µM CP-91149 for 1.5 hours. Glycogen contents were determined and normalized to protein content. Data are expressed as % of the baseline value (224 ± 16 nmol glucosyl unit per mg protein, n = 3) and mean ± SD. (F) H69 cholangiocytes were treated as in (B) and glycogen content of the cells was determined. Data are expressed as % of the baseline value (52 ± 3 nmol glucosyl unit per mg protein, n = 3) and mean ± SD. (G) H69 cholangiocytes were preincubated with 5.5 mM glucose for 1 hour. After harvesting baseline samples, cells were treated with 0.1% DMSO (vehicle control), 50 µM LRE1, or 1 µM Forskolin for 1 hour. Glycogen contents were determined and normalized to protein content. Data are expressed as % of the baseline value (213 ± 8 nmol glucosyl unit per mg protein, n = 3) and mean ± SD. All data are representative of 2-3 independent experiments.

### sAC- and tmAC-derived cAMP have different effectors to regulate glycogen homeostasis

We next investigated by which effector sAC-derived cAMP exerts its effect on glycogen. sAC-derived cAMP has been shown to signal via both Epac (Flacke et al., 2013; Onodera et al., 2014) and PKA (Acin-Perez et al., 2009; Valsecchi et al., 2017). In glucose-starved HepG2 cells that were fuelled with only octanoate, inhibition of Epac1 by the specific inhibitor (R)-CE3F4 (Courilleau et al., 2012) induced an acute, transient increase in ECAR with the same temporal and dynamic characteristics as sAC inhibition (Figure 3A). In contrast, the PKA inhibitor H89 was without any effect (Figure 3A). The Epac2-specific inhibitor ESI-05 (Tsalkova et al., 2012) also did not induce changes in ECAR (data not shown).

**Figure 3.**
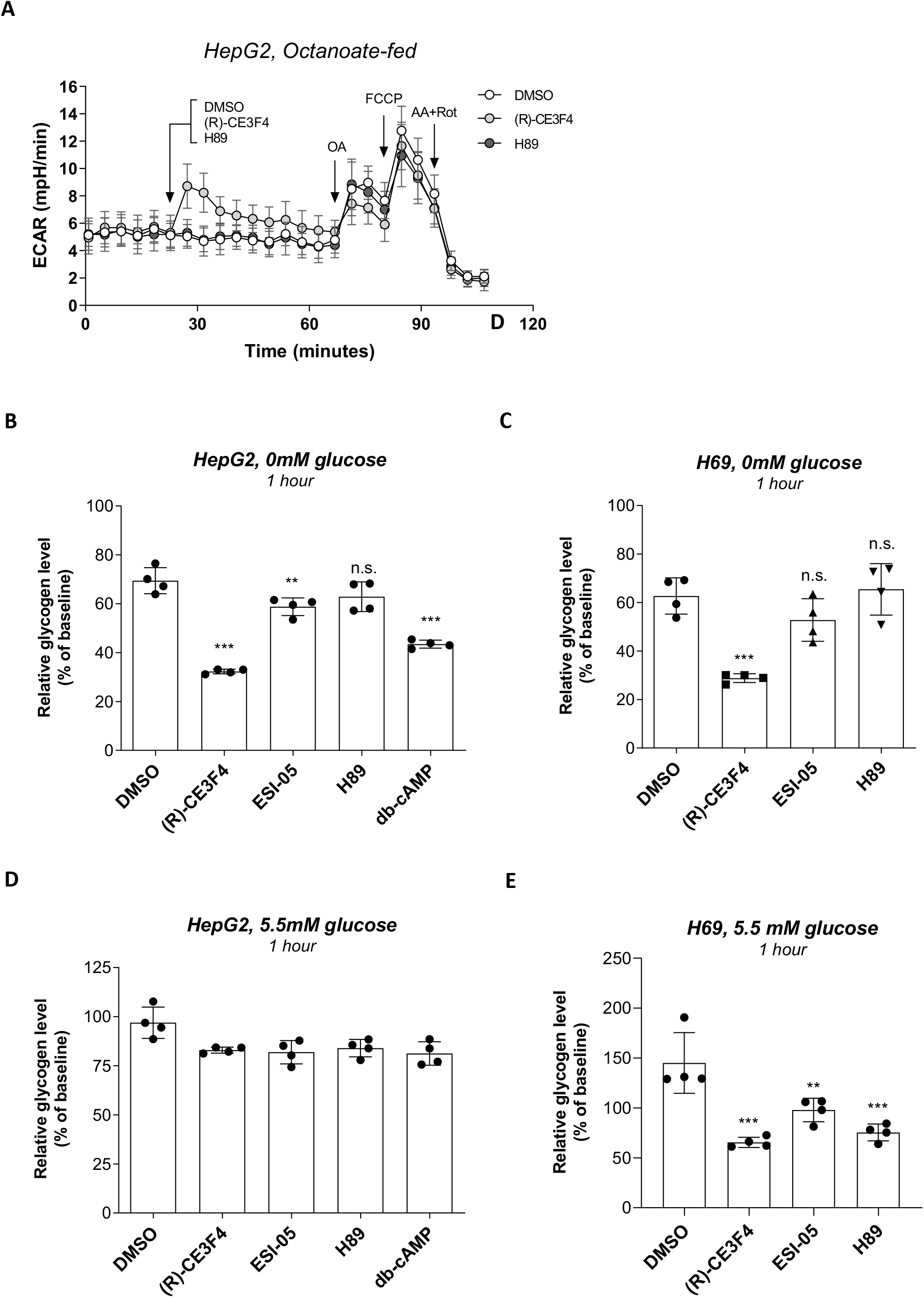
sAC- and tmAC-derived cAMP have different effectors to regulate glycogen homeostasis. (A) HepG2 cells were pre-incubated for 1 hour in HBSS for ambient air with 125 µM octanoate and then the extracellular acidification rate (ECAR) was measured with the Seahorse Flux Analyzer XF96. After baseline measurement, 0.1% DMSO (vehicle control, n=6), 50 µM (R)-CE3F4 (n=6), or 10 µM H89 (n=5) was injected, followed by oligomycin A (OA), carbonyl cyanide-p-trifluoromethoxy-phenyl hydrazone (FCCP), and antimycin A (AA) and rotenone (Rot). Data are presented as mean ± SD. (B-C) HepG2 cells (B) and H69 cholangiocytes (C) were acutely incubated in glucose-free medium for 60 min in the presence of 0.1% DMSO (vehicle control), 50 µM (R)-CE3F4, 10 µM ESI-05, or 10 µM H89. HepG2 cells were also treated with 100 µM dibutyryl-cAMP (db-cAMP) as positive control. Glycogen content was determined and normalized to protein content. Data are expressed as % of the baseline value (HepG2: 71 ± 3 nmol glucosyl unit per mg protein, n = 4. H69: 130 ± 13 nmol glucosyl unit per mg protein, n = 4) and mean ± SD. (D-E) HepG2 cells (D) and H69 cholangiocytes (E) were incubated in medium containing 5.5 mM glucose for 1 hour in the presence of 0.1% DMSO (vehicle control), 50 µM (R)-CE3F4, 10 µM ESI-05, or 10 µM H89. HepG2 cells were also treated with 100 µM dibutyryl-cAMP (db-cAMP) as positive control. Glycogen content was determined and normalized to protein content. Data are expressed as % of the baseline value (HepG2: 74 ± 2 nmol glucosyl unit per mg protein, n = 4. H69: 147 ± 9 nmol glucosyl unit per mg protein, n = 4) and mean ± SD. All data are representative of 2-3 independent experiments.

To corroborate the observed ECAR changes, we examined the effects of inhibition of Epac1, Epac2, and PKA on cellular glycogen levels. Indeed, in HepG2 cells acutely deprived of glucose, only inhibition of Epac1 phenocopied the glycogenolysis induced by sAC inhibition (Figure 3B). In addition, the PKA-selective activator dibutyryl-cAMP promoted glycogen breakdown, confirming that PKA and Epac1 mediate opposite metabolic effects of tmAC and sAC signalling, respectively. A similar result was obtained in H69 cells incubated without glucose (Figure 3C). Moreover, we observed that in the presence of glucose, the Epac1-specific inhibitor (R)-CE3F4 induced significant glycogen breakdown in H69 but not in HepG2 cells (Figure 3D and 3E), which mirrored the differential effects of sAC inhibition on glycogen in these cells. Of note, inhibition of sAC or Epac1 also induced glycogenolysis and the concomitant release of lactate in several other cell lines (Figure S3A-S3D). Taken together, our data suggest that Epac1 mediates the sAC-dependent repression of glycogenolysis in several cell types, whereas tmAC-activated PKA oppositely induces glycogenolysis.

### Glucose deprivation reveals a complex I-independent regulation of glycogenolysis by sAC

We and others have shown that sAC regulates the activity of complex I of the respiratory chain (Chang et al., 2021; De Rasmo et al., 2015; Valsecchi et al., 2017). Inhibition of complex I can reduce the steady state glycogen level via AMPK-dependent suppression of glycogen synthase activity (Bultot et al., 2012; Carling and Hardie, 1989; Heinz et al., 2017). Indeed, in cultured primary murine astrocytes, sAC inhibition suppresses oxidative phosphorylation, leading to AMPK activation and a decrease in glycogen level (Jakobsen et al., 2021). However, because sAC inhibition also induces glycogen breakdown in the absence glucose (Figure 1 and 2), leaving glycogen synthase without substrate, we examined whether the induction of glycogenolysis upon sAC inhibition was mediated by complex I suppression or by another, as yet unknown, mechanism. To this end, we examined whether the complex I inhibitor rotenone could phenocopy the metabolic effects of sAC inhibition. In glucose-fed HepG2 cells, 15 nM rotenone suppressed both coupled and uncoupled OCR to a comparable extent as 50 µM LRE1 (Figure 4A). However, in the absence of glucose, rotenone caused a much smaller ECAR transient than LRE1 despite comparable inhibition of complex I activity (Figure 4B). Rotenone had also a much smaller effect on lactate release under these conditions (Figure 4C). Although the addition of rotenone to HepG2 cells did result in clear AMPK activation even to a greater extent than LRE1 did (Figure S4A), this dose of rotenone did not induce significant glycogen breakdown as indicated by the inability to induce clear ECAR changes. Thus, our data obtained under conditions of glucose deprivation, strongly suggest that sAC also regulates glycogenolysis independent of its regulation of complex I.

**Figure 4.**
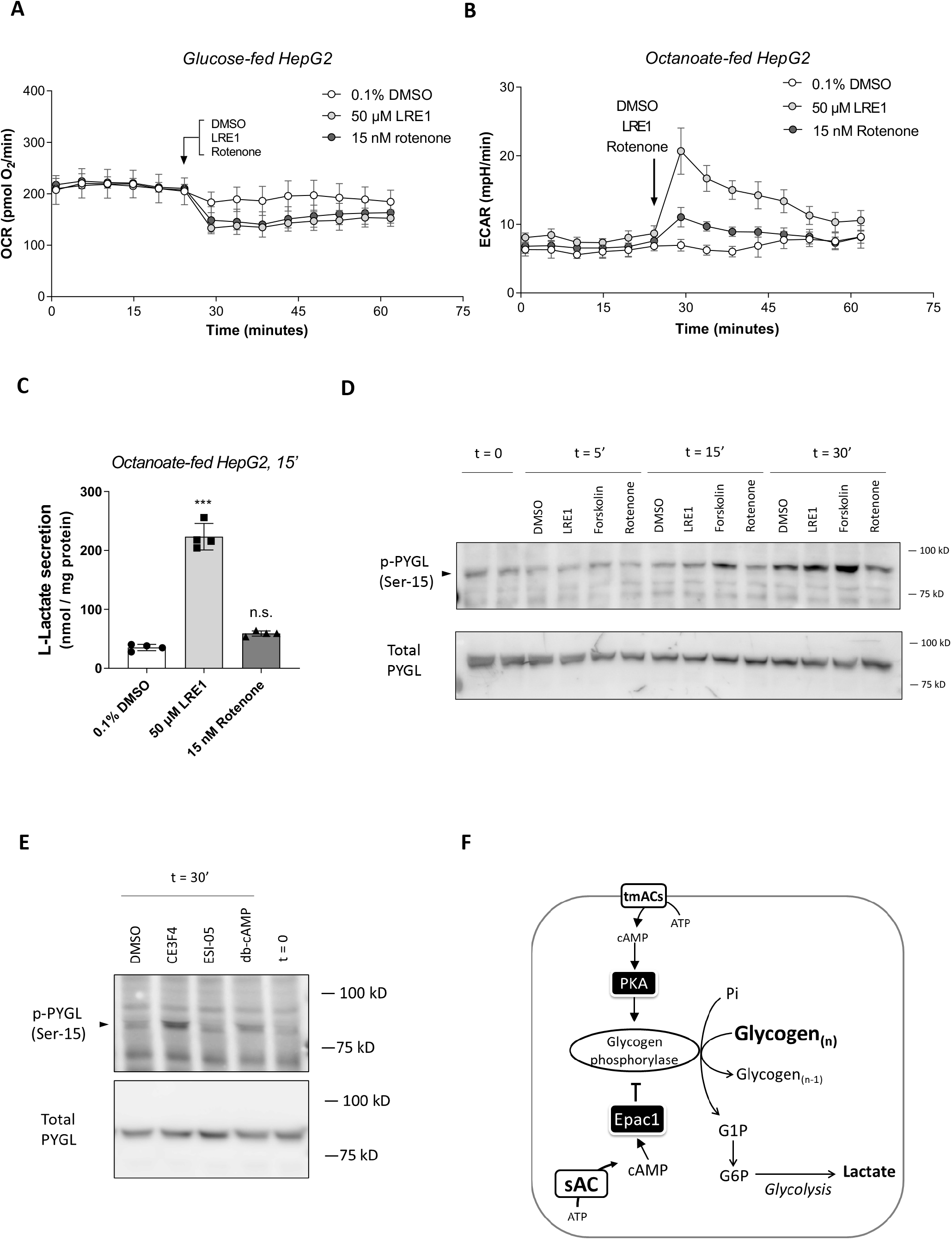
The effects of partial inhibition of complex I on glycogenolysis. (A) HepG2 cells were pre-incubated in HBSS for ambient air in the presence of 5.5 mM glucose for 30 minutes and then the oxygen consumption rate (OCR) was measured with the Seahorse Flux Analyzer XF96. After baseline measurement, 0.1% DMSO (vehicle control, n=8), 50 µM LRE1 (n=5) or 15 nM rotenone (n=6) was injected. (B) HepG2 cells were pre-incubated in HBSS for ambient air in the presence of 125 µM octanoate for 30 min and then the oxygen consumption rate (ECAR) was measured with the Seahorse Flux Analyzer XF96. After baseline measurement, 0.1% DMSO (vehicle control, n=6), 50 µM LRE1 (n=6) or 15 nM rotenone (n=6) was injected. (C) HepG2 cells were pre-incubated in glucose-free HBSS with 125 µM octanoate for 30 min and then the medium was replaced by the same incubation medium containing 0.1% DMSO (vehicle control), 50 µM LRE1 or 15 nM rotenone. After 15 min of incubation, samples were taken from the extracellular medium to determine lactate release. Data are presented as mean ± SD of quadruplicate determination. (D) HepG2 cells were incubated in serum-free DMEM containing 5.5 mM glucose for 1 hour. Cells were then treated with 0.1% DMSO (vehicle control), 50 µM LRE1, 1 µM forskolin or 15 nM rotenone for 5, 15 and 30 minutes. Phosphorylation of Ser-15 of liver form glycogen phosphorylase (PYGL) was examined by immunoblotting. (E) HepG2 cells were treated as in (D) but treated with 50 µM (R)-CE3F4, 10 µM ESI-05, or 100 µM db-cAMP for 30 min. Phosphorylation of Ser-15 of liver form glycogen phosphorylase (PYGL) was examined by immunoblotting. (F) Schematic representation of key findings of this study. Data are representative of 2-3 independent experiments.

### Activation of glycogen phosphorylase a by sAC-cAMP-Epac1 signalling

We next sought to characterize how sAC regulates glycogenolysis independently of complex I. The glycogen content of cells depends on the equilibrium between of glycogen synthase and glycogen phosphorylase activity. Since the inhibition of sAC-cAMP-Epac1 signalling promoted glycogenolysis in the absence of glucose, leaving glycogen synthase without substrate, we anticipated that inhibition of sAC-cAMP-Epac1 signalling would increase glycogen phosphorylase activity. The conversion of glycogen phosphorylase from the inactive T state to the active R state is stimulated allosterically by AMP and covalently by phosphorylase kinase-mediated phosphorylation at residue Ser-15 (Johnson, 1992). Since sAC inhibition did not cause changes in total ATP production, the adenylate energy charge, or the level of AMP (data not shown) both in the absence (Figure S1E and S1F) and presence (Chang et al., 2021) of glucose, we hypothesized that sAC regulates the phosphorylation state of glycogen phosphorylase. To avoid inadvertent activation of AMPK during glucose deprivation, we performed these experiments in the presence of a physiological concentration of glucose (5.5 mM). Indeed, both the sAC-specific inhibitor LRE1 as well as the tmAC-specific activator forskolin increased Ser-15 phosphorylation of glycogen phosphorylase (PYGL) (Figure 4D). Similarly, both the Epac1-selective inhibitor (R)-CE3F4 and the PKA-selective activator dibutyryl-cAMP induced phosphorylation of Ser-15 in PYGL, whereas 15 nM rotenone was without effect (Figure 4D). Similar results on glycogen phosphorylase phosphorylation were obtained in H69 human cholangiocytes (Figure S4B and S4C).

In conclusion, our data show that sAC-derived cAMP and tmAC-derived cAMP function within independent signalling microdomains that have opposite effects on glycogenolysis: sAC-cAMP-Epac1 signalling suppresses glycogenolysis while tmAC-cAMP-PKA signalling promotes glycogenolysis. While the hormone-stimulated tmAC-cAMP-PKA mediates glycogenolysis in cells specialized in glycogen storage, we suggest that sAC-cAMP-Epac1 signalling regulates glycogenolysis independent of hormonal signalling to maintain energy homeostasis in both specialized and non-specialized cells. From the viewpoint of autonomous cellular metabolism, the cellular ATP level feeds back to the sAC-cAMP-Epac1 signalling to prevent unnecessary glycogenolysis and wasteful lactate secretion during glucose starvation and, in some cell types, also in the presence of glucose. Our data strongly underscore the versatility of cAMP signalling and the concept of signalling microdomains, where adenylyl cyclases, cAMP-degrading phosphodiesterases, cAMP effectors, and downstream substrates confine cAMP signalling to specific locations within cells (Kamenetsky et al., 2006; Lefkimmiatis and Zaccolo, 2014).

## Supporting information

Method and Legends for Supplementary Figures

## Acknowledgement

The authors thank Dr. de Korte and his colleagues (Sanquin Blood Supply Foundation, Amsterdam, NL) for performing the HPLC analysis of cytosolic adenine nucleotides. JC is supported by the Talent Development Grant from the Amsterdam Gastroenterology, Endocrinology & Metabolism (AGEM) Research Institute. PB is supported by the Brazilian National Council for Scientific and Technological Development — CNPq.

## Author Contributions

AV performed experiments and wrote the paper, PB, GH, SG and HL performed experiments, ROE supervised the study and JC designed the study and wrote the paper.

## Legends to the Figures

**Supplemental Figure S1.**
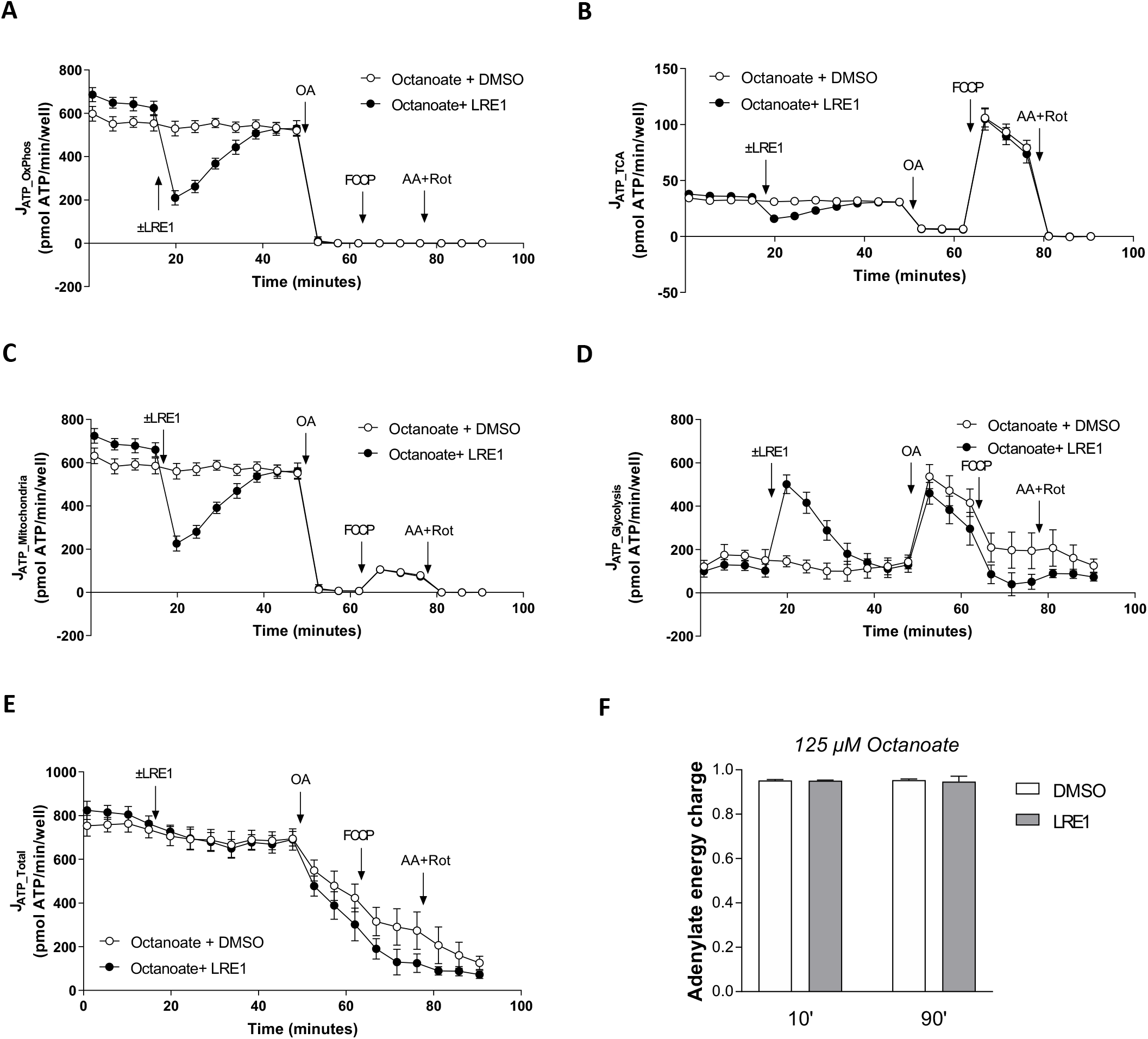

**Supplemental Figure S2.**
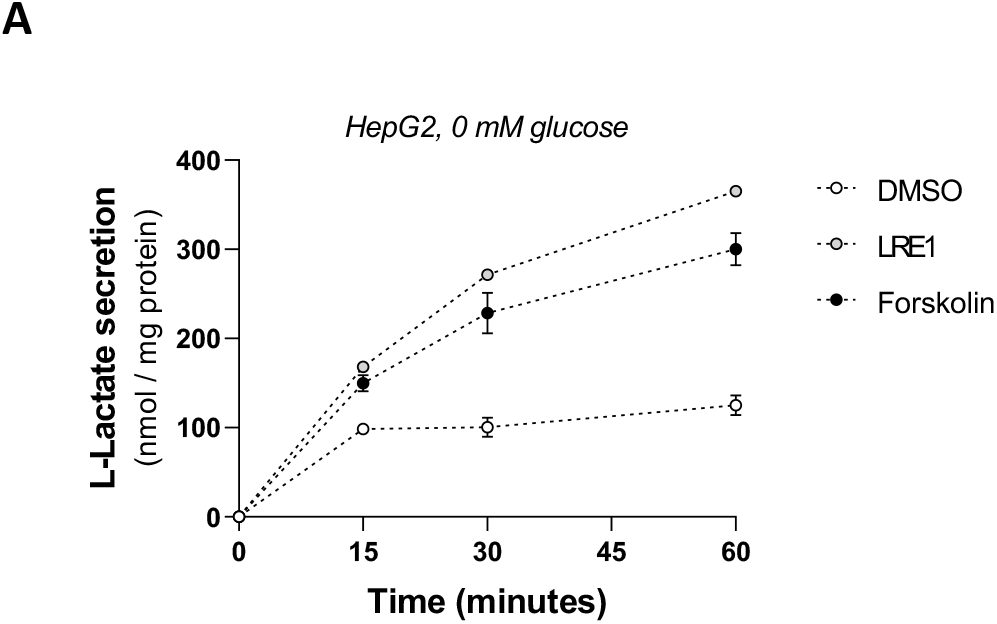

**Supplemental Figure S3.**
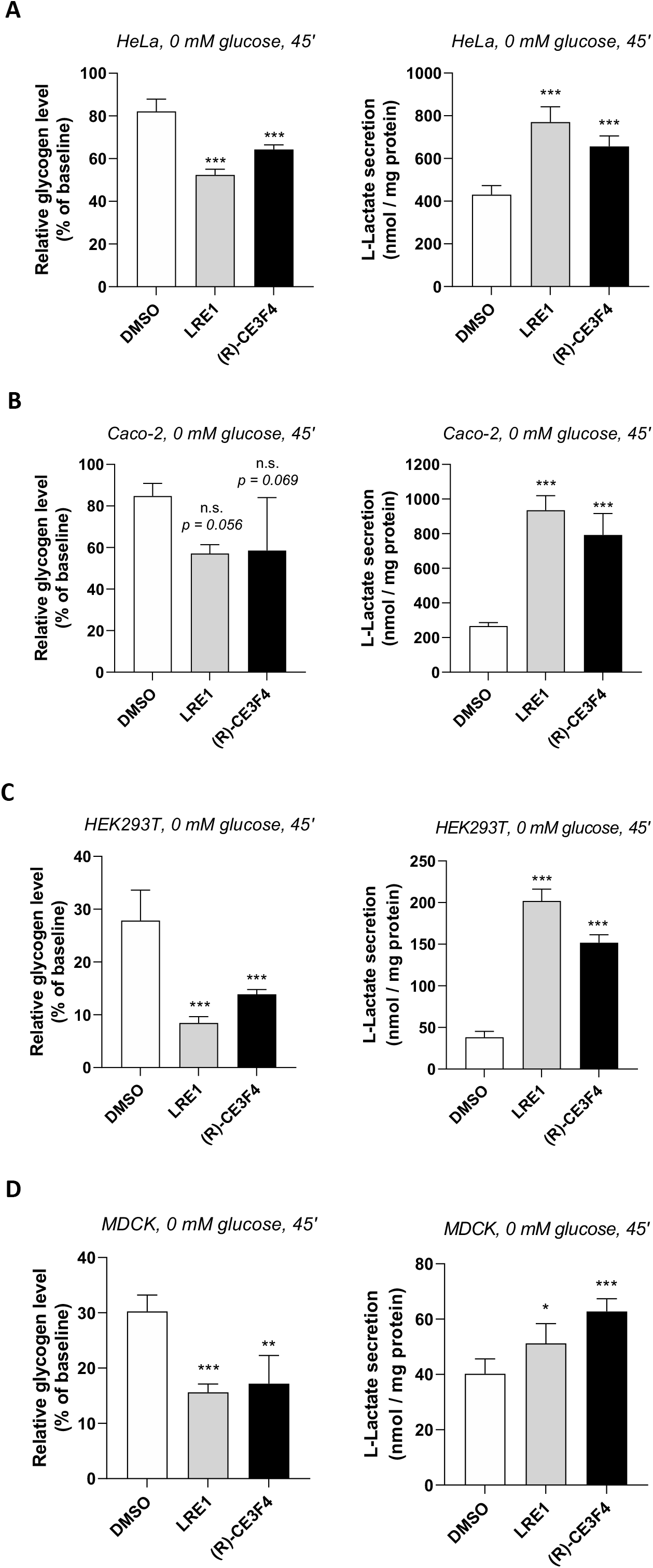

**Supplemental Figure S4.**
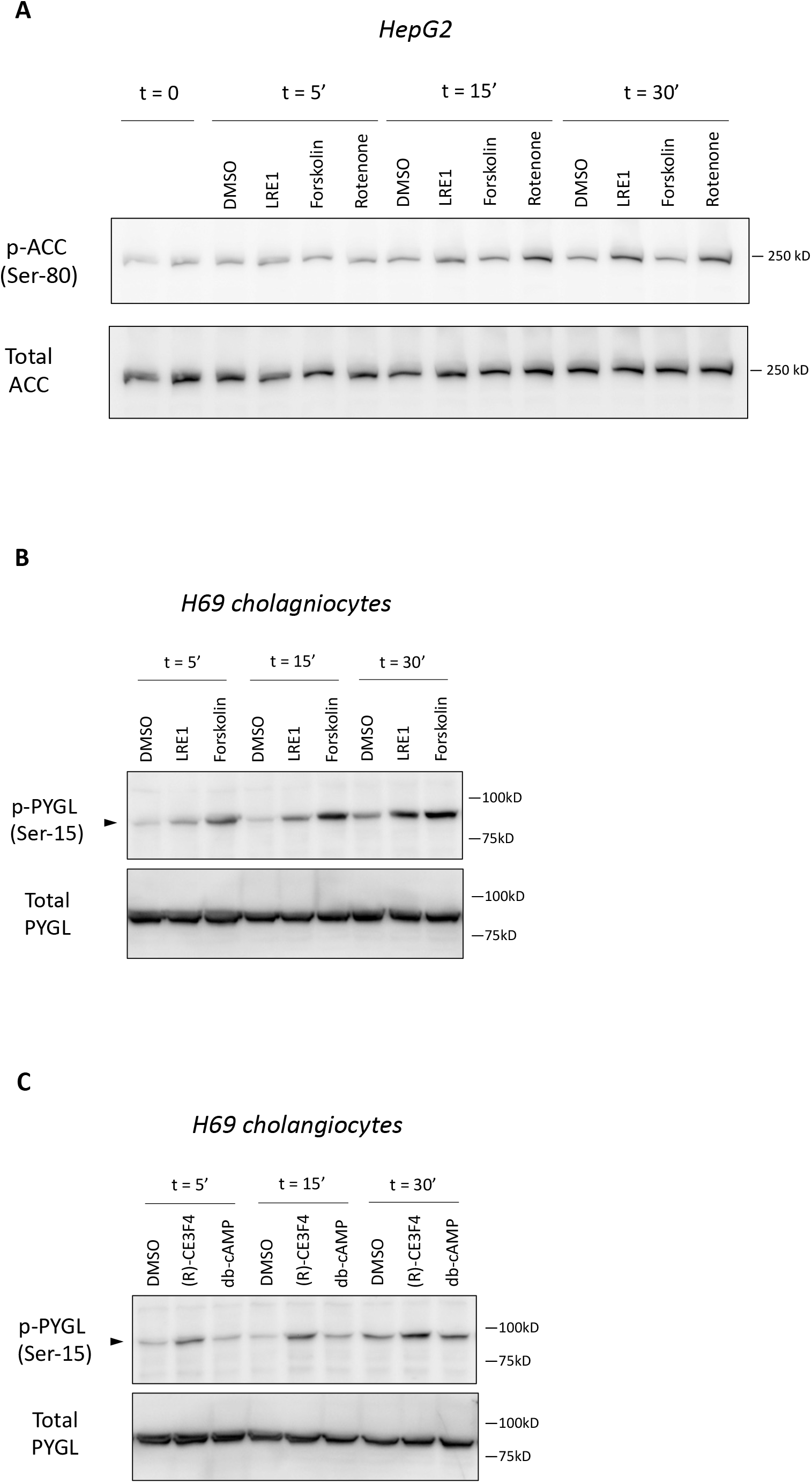

## References

Acin-Perez, R., Salazar, E., Kamenetsky, M., Buck, J., Levin, L.R., and Manfredi, G. (2009). Cyclic AMP produced inside mitochondria regulates oxidative phosphorylation. Cell Metab 9, 265–276.

Agarwal, S.R., Clancy, C.E., and Harvey, R.D. (2016). Mechanisms Restricting Diffusion of Intracellular cAMP. Sci Rep 6, 19577.

Agarwal, S.R., Sherpa, R.T., Moshal, K.S., and Harvey, R.D. (2022). Compartmentalized cAMP signaling in cardiac ventricular myocytes. Cell Signal 89, 110172.

Berthet, J., Rall, T.W., and Sutherland, E.W. (1957). The relationship of epinephrine and glucagon to liver phosphorylase. IV. Effect of epinephrine and glucagon on the reactivation of phosphorylase in liver homogenates. J Biol Chem 224, 463–475.

Buck, J., Sinclair, M.L., Schapal, L., Cann, M.J., and Levin, L.R. (1999). Cytosolic adenylyl cyclase defines a unique signaling molecule in mammals. Proc Natl Acad Sci U S A 96, 79–84.

Bultot, L., Guigas, B., Von Wilamowitz-Moellendorff, A., Maisin, L., Vertommen, D., Hussain, N., Beullens, M., Guinovart, J.J., Foretz, M., Viollet, B., et al. (2012). AMP-activated protein kinase phosphorylates and inactivates liver glycogen synthase. Biochem J 443, 193–203.

Carling, D., and Hardie, D.G. (1989). The substrate and sequence specificity of the AMP-activated protein kinase. Phosphorylation of glycogen synthase and phosphorylase kinase. Biochim Biophys Acta 1012, 81–86.

Chang, J.C., Go, S., de Waart, D.R., Munoz-Garrido, P., Beuers, U., Paulusma, C.C., and Oude Elferink, R. (2016). Soluble Adenylyl Cyclase Regulates Bile Salt-Induced Apoptosis in Human Cholangiocytes. Hepatology 64, 522–534.

Chang, J.C., Go, S., Gilglioni, E.H., Duijst, S., Panneman, D.M., Rodenburg, R.J., Li, H.L., Huang, H.L., Levin, L.R., Buck, J., et al. (2021). Soluble adenylyl cyclase regulates the cytosolic NADH/NAD(+) redox state and the bioenergetic switch between glycolysis and oxidative phosphorylation. Biochim Biophys Acta Bioenerg 1862, 148367.

Chen, Y., Cann, M.J., Litvin, T.N., Iourgenko, V., Sinclair, M.L., Levin, L.R., and Buck, J. (2000). Soluble adenylyl cyclase as an evolutionarily conserved bicarbonate sensor. Science 289, 625–628.

Choi, H.B., Gordon, G.R., Zhou, N., Tai, C., Rungta, R.L., Martinez, J., Milner, T.A., Ryu, J.K., McLarnon, J.G., Tresguerres, M., et al. (2012). Metabolic communication between astrocytes and neurons via bicarbonate-responsive soluble adenylyl cyclase. Neuron 75, 1094–1104.

Courilleau, D., Bisserier, M., Jullian, J.C., Lucas, A., Bouyssou, P., Fischmeister, R., Blondeau, J.P., and Lezoualc’h, F. (2012). Identification of a tetrahydroquinoline analog as a pharmacological inhibitor of the cAMP-binding protein Epac. J Biol Chem 287, 44192–44202.

De Rasmo, D., Signorile, A., Santeramo, A., Larizza, M., Lattanzio, P., Capitanio, G., and Papa, S. (2015). Intramitochondrial adenylyl cyclase controls the turnover of nuclear-encoded subunits and activity of mammalian complex I of the respiratory chain. Biochim Biophys Acta 1853, 183–191.

Di Benedetto, G., Scalzotto, E., Mongillo, M., and Pozzan, T. (2013). Mitochondrial Ca(2)(+) uptake induces cyclic AMP generation in the matrix and modulates organelle ATP levels. Cell Metab 17, 965–975.

Fischer, E.H., and Krebs, E.G. (1955). Conversion of phosphorylase b to phosphorylase a in muscle extracts. J Biol Chem 216, 121–132.

Flacke, J.P., Flacke, H., Appukuttan, A., Palisaar, R.J., Noldus, J., Robinson, B.D., Reusch, H.P., Zippin, J.H., and Ladilov, Y. (2013). Type 10 soluble adenylyl cyclase is overexpressed in prostate carcinoma and controls proliferation of prostate cancer cells. J Biol Chem 288, 3126–3135.

Fredriksson, R., and Schioth, H.B. (2005). The repertoire of G-protein-coupled receptors in fully sequenced genomes. Mol Pharmacol 67, 1414–1425.

Geng, W., Wang, Z., Zhang, J., Reed, B.Y., Pak, C.Y., and Moe, O.W. (2005). Cloning and characterization of the human soluble adenylyl cyclase. Am. J. Physiol Cell Physiol 288, C1305–C1316.

Hanoune, J., and Defer, N. (2001). Regulation and role of adenylyl cyclase isoforms. Annu Rev Pharmacol Toxicol 41, 145–174.

Heinz, S., Freyberger, A., Lawrenz, B., Schladt, L., Schmuck, G., and Ellinger-Ziegelbauer, H. (2017). Mechanistic Investigations of the Mitochondrial Complex I Inhibitor Rotenone in the Context of Pharmacological and Safety Evaluation. Sci Rep 7, 45465.

Jaiswal, B.S., and Conti, M. (2003). Calcium regulation of the soluble adenylyl cyclase expressed in mammalian spermatozoa. Proc. Natl. Acad. Sci. U. S. A 100, 10676–10681.

Jakobsen, E., Andersen, J.V., Christensen, S.K., Siamka, O., Larsen, M.R., Waagepetersen, H.S., Aldana, B.I., and Bak, L.K. (2021). Pharmacological inhibition of mitochondrial soluble adenylyl cyclase in astrocytes causes activation of AMP-activated protein kinase and induces breakdown of glycogen. Glia 69, 2828–2844.

Johnson, L.N. (1992). Glycogen phosphorylase: control by phosphorylation and allosteric effectors. FASEB J 6, 2274–2282.

Jurevicius, J., and Fischmeister, R. (1996). cAMP compartmentation is responsible for a local activation of cardiac Ca2+ channels by beta-adrenergic agonists. Proc Natl Acad Sci U S A 93, 295–299.

Kamenetsky, M., Middelhaufe, S., Bank, E.M., Levin, L.R., Buck, J., and Steegborn, C. (2006). Molecular details of cAMP generation in mammalian cells: a tale of two systems. J Mol Biol 362, 623–639.

Kleinboelting, S., Diaz, A., Moniot, S., van den Heuvel, J., Weyand, M., Levin, L.R., Buck, J., and Steegborn, C. (2014). Crystal structures of human soluble adenylyl cyclase reveal mechanisms of catalysis and of its activation through bicarbonate. Proc Natl Acad Sci U S A 111, 3727–3732.

Kobayashi, M., Buck, J., and Levin, L.R. (2004). Conservation of functional domain structure in bicarbonate-regulated “soluble” adenylyl cyclases in bacteria and eukaryotes. Dev Genes Evol 214, 503–509.

Kumar, S., Kostin, S., Flacke, J.P., Reusch, H.P., and Ladilov, Y. (2009). Soluble adenylyl cyclase controls mitochondria-dependent apoptosis in coronary endothelial cells. J Biol Chem 284, 14760–14768.

Lefkimmiatis, K., and Zaccolo, M. (2014). cAMP signaling in subcellular compartments. Pharmacol Ther 143, 295–304.

Levin, L.R., and Buck, J. (2015). Physiological roles of acid-base sensors. Annu Rev Physiol 77, 347–362.

Litvin, T.N., Kamenetsky, M., Zarifyan, A., Buck, J., and Levin, L.R. (2003). Kinetic properties of “soluble” adenylyl cyclase. Synergism between calcium and bicarbonate. J. Biol. Chem 278, 15922–15926.

Martin, W.H., Hoover, D.J., Armento, S.J., Stock, I.A., McPherson, R.K., Danley, D.E., Stevenson, R.W., Barrett, E.J., and Treadway, J.L. (1998). Discovery of a human liver glycogen phosphorylase inhibitor that lowers blood glucose in vivo. Proc Natl Acad Sci U S A 95, 1776–1781.

Mookerjee, S.A., Gerencser, A.A., Nicholls, D.G., and Brand, M.D. (2017). Quantifying intracellular rates of glycolytic and oxidative ATP production and consumption using extracellular flux measurements. J Biol Chem 292, 7189–7207.

Mookerjee, S.A., Goncalves, R.L., Gerencser, A.A., Nicholls, D.G., and Brand, M.D. (2015). The contributions of respiration and glycolysis to extracellular acid production. Biochim Biophys Acta 1847, 171–181.

Obiako, B., Calchary, W., Xu, N., Kunstadt, R., Richardson, B., Nix, J., and Sayner, S.L. (2013). Bicarbonate disruption of the pulmonary endothelial barrier via activation of endogenous soluble adenylyl cyclase, isoform 10. Am J Physiol Lung Cell Mol Physiol 305, L185–192.

Onodera, Y., Nam, J.M., and Bissell, M.J. (2014). Increased sugar uptake promotes oncogenesis via EPAC/RAP1 and O-GlcNAc pathways. J Clin Invest 124, 367–384.

Pidoux, G., and Tasken, K. (2010). Specificity and spatial dynamics of protein kinase A signaling organized by A-kinase-anchoring proteins. J Mol Endocrinol 44, 271–284.

Ramos-Espiritu, L., Kleinboelting, S., Navarrete, F.A., Alvau, A., Visconti, P.E., Valsecchi, F., Starkov, A., Manfredi, G., Buck, H., Adura, C., et al. (2016). Discovery of LRE1 as a specific and allosteric inhibitor of soluble adenylyl cyclase. Nat Chem Biol 12, 838–844.

Rousset, M., Zweibaum, A., and Fogh, J. (1981). Presence of glycogen and growth-related variations in 58 cultured human tumor cell lines of various tissue origins. Cancer Res 41, 1165–1170.

Sayner, S.L., Frank, D.W., King, J., Chen, H., VandeWaa, J., and Stevens, T. (2004). Paradoxical cAMP-induced lung endothelial hyperpermeability revealed by Pseudomonas aeruginosa ExoY. Circ Res 95, 196–203.

Sutherland, E.W., and Rall, T.W. (1957). THE PROPERTIES OF AN ADENINE RIBONUCLEOTIDE PRODUCED WITH CELLULAR PARTICLES, ATP, Mg++, AND EPINEPHRINE OR GLUCAGON. Journal of the American Chemical Society 79, 3608–3608.

Sutherland, E.W., and Rall, T.W. (1958). Fractionation and characterization of a cyclic adenine ribonucleotide formed by tissue particles. J Biol Chem 232, 1077–1091.

Tsalkova, T., Mei, F.C., Li, S., Chepurny, O.G., Leech, C.A., Liu, T., Holz, G.G., Woods, V.L., Jr., and Cheng, X. (2012). Isoform-specific antagonists of exchange proteins directly activated by cAMP. Proc Natl Acad Sci U S A 109, 18613–18618.

Valsecchi, F., Konrad, C., D’Aurelio, M., Ramos-Espiritu, L.S., Stepanova, A., Burstein, S.R., Galkin, A., Magrane, J., Starkov, A., Buck, J., et al. (2017). Distinct intracellular sAC-cAMP domains regulate ER Ca(2+) signaling and OXPHOS function. J Cell Sci 130, 3713–3727.

Walsh, D.A., Perkins, J.P., and Krebs, E.G. (1968). An adenosine 3’,5’-monophosphate-dependant protein kinase from rabbit skeletal muscle. J Biol Chem 243, 3763–3765.

Zaccolo, M., and Pozzan, T. (2002). Discrete microdomains with high concentration of cAMP in stimulated rat neonatal cardiac myocytes. Science 295, 1711–1715.

Zippin, J.H., Chen, Y., Straub, S.G., Hess, K.C., Diaz, A., Lee, D., Tso, P., Holz, G.G., Sharp, G.W., Levin, L.R., et al. (2013). CO2/HCO3(-)- and calcium-regulated soluble adenylyl cyclase as a physiological ATP sensor. J Biol Chem 288, 33283–33291.

