## Supplementary material for "Opposite regulation of glycogen metabolism by cAMP produced in the cytosol and at the plasma membrane": Method and Legends for Supplementary Figures

### Supplemental Information

#### METHODS

##### Materials

Unless otherwise indicated, materials were purchased from Sigma-Aldrich.

##### Cell culture

The immortalized human intrahepatic cholangiocyte H69 cell line was a kind gift from Douglas Jefferson (Grubman et al., 1994) and was cultured in DMEM/F-12 (3:1) with hormonal supplements as described (Chang et al., 2016). HepG2, Caco-2, HeLa, HEK293T, Madin-Darby canine kidney (MDCK) cells were cultured in DMEM (Invitrogen), supplemented with 2 g/L glucose, 1.8 g/L NaHCO3, 20 mM HEPES-NaOH pH 7.4 and 10% fetal bovine serum. For all experiments, cells were refreshed the day before the experiment.

##### Extracellular flux analysis

Extracellular acidification rate (ECAR) and oxygen consumption rate (OCR) were measured with a Seahorse XF96 Analyzer (Agilent, United States). HepG2 cells were cultured in a 96-well Seahorse culture plate until confluence. Media were refreshed the day before the experiment. On the day of the experiment, cells were incubated for 1 hour at 37°C in experimental medium, which was based on modified Hank's balanced salt solution (HBSS) for ambient air (please refer to Table S1 for formulation) supplemented with 0.1% fatty acid-free bovine serum albumin (BSA). We did not observe an effect of amino acids on the acute regulation of glycogen metabolism by sAC. During this pre-incubation the indicated substrates or inhibitors (2-deoxyglucose and glycogen phosphorylase a inhibitor CP-91149) were added. Twenty-five microliters of concentrated compound solutions prepared in experimental medium (without BSA) were injected as indicated. The final concentrations of compounds were as follows: LRE1, 50 µM; oligomycin A, 2.5 µM; FCCP, 2 µM; antimycin A, 2.5 µM; rotenone, 1 µM. In the experiments of Figures 3 and 4, LRE1 was replaced by the inhibitors indicated in the figure. The coupled respiration rate was defined as the difference between the average of the last three OCR measurements after the addition of inhibitors or vehicle control and the last OCR measurement after the addition of oligomycin A. The FCCP-driven respiration rate was defined as the difference between the first OCR measurement after FCCP addition and the last OCR measurement after the addition of oligomycin A.

**Estimation of ATP production by glycolysis and by mitochondria using extracellular flux measurements**

The time-lapsed ATP production rates were calculated from extracellular fluxes (ECARs and OCRs) as elegantly described (Mookerjee et al., 2017; Mookerjee et al., 2015). Since cells were pre-incubated in HBSS until reaching steady state and before imposing the acute perturbation of sAC activity, it was assumed that the oxidation of the provided substrates in the mitochondria was complete and that the contribution from other endogenous substrates (except for glycogen) was negligible. Because cells were fueled solely with octanoate, the glycolytic flux was assumed to come from glycogenolysis. The total ATP production rate ($J_{ATP\_total}$) is defined as the sum of the ATP production rate of glycolysis ($J_{ATP\_glycolysis}$, substrate level phosphorylation by phosphoglycerate kinase and pyruvate kinase) and the ATP production rate of mitochondria ($J_{ATP\_mitochondria}$, substrate level phosphorylation by succinyl-CoA synthetase in the TCA cycle and oxidative phosphorylation by the electron transport chain and ATP synthase). Further details of the calculation of ATP production rates are given in (Chang et al., 2021). The values of maximal $H^+/O_2$, ATP/ lactate, maximal P/O ratios with octanoate and glycogen as bioenergetic substrates are given in Table S2.

**Sample preparation for the enzymatic determination of glycogen**

At the end of the incubation, cells were washed twice with ice-cold PBS and lysed in TTE buffer (1% Triton X-100, 10 mM Tris-HCl pH 8.0 and 1 mM EDTA-NaOH, pH 8.0) or RIPA buffer (150 mM NaCl, 20 mM Tris-HCl pH 8.0, 1% Triton X-100, 0.1% SDS, 0.5% Na-deoxycholate). Lysates were centrifuged at 20,000 × g at 4°C for 10 min. Supernatants were harvested for determination of glycogen and protein concentration. For the determination of glycogen, 50 µL of supernatant was mixed with 20 µL 0.35 M NaOH and heated at 80°C for 30 minutes to degrade monosaccharides. The hot alkali-treated lysates were then deproteinized by adding 30 µL 10% (w/v) metaphosphoric acid (MPA) and incubated on ice for at least 1 hour or overnight at 4°C. Samples were centrifuged at 20,000 × g for 10 min. Twenty microliters of deproteinized samples or glucose standards (prepared in 3% MPA) were mixed with 100 µL solution A (2.5 U/mL amyloglucosidase from *Aspergillus niger*, 50 mM $K_2HPO_4$-$KH_2PO_4$, pH 8.0). The resulting mixture had a pH of about 4.7, which is optimal for amyloglucosidase to hydrolyze glycogen. After 1-hour incubation at 45°C, 50 µL solution B was added (1.5 mM homovanillic acid, 2 U/mL horseradish peroxidase, 0.5 M $K_2HPO_4$-$KH_2PO4$, pH 8.0) to correct the pH to 6.8 to enable the measurement of the glucose produced by glucose oxidase. After determining the background fluorescence at $\lambda_{ex}/\lambda_{em}$= 320/450 nm in the CLARIOstar microplate reader (BMG LABTECH, Ortenberg, Germany), 50 µL start solution (containing 8 U/mL glucose oxidase) was added and fluorescence was followed every 2 minutes until the reaction was complete (within 1 hour).

**Isolation and culture of primary mouse hepatocytes**

Animal experiments were approved by the institutional animal experiment committee. Primary mouse hepatocytes were isolated from wild-type male C57BL/6J mice after overnight *ad libitum* feeding by a two-step collagenase perfusion method through the portal vein. Cells were cultured in collagen sandwich configuration overnight on a 60-rpm shaking platform in a 5% $CO_2$, 37°C incubator. The isolation procedure and culture of primary mouse hepatocytes were performed as described (Gilglioni et al., 2018).

**Sample preparation to measure lactate secretion**

Cells were refreshed with full culture medium one day prior to the experiment. On the day of the experiment, medium was changed to the experimental medium (for composition see above) supplemented with 5.5 mM glucose and 0.1% fatty acid-free bovine serum albumin (BSA). At the start of the experiment, cells were exposed to media containing the indicated inhibitors and vehicle controls. At indicated time points, 50 µL of spent medium was mixed with 75 µL ice-cold 5% (w/v) meta-phosphoric acid (MPA) for deproteinization. After incubating for at least one hour at 4°C, the MPA-acidified samples were centrifuged at 20,000 × *g* for 10 min. The supernatants were harvested for enzymatic determination of L-lactate using the LDH-catalysed reduction of $NAD^+$ as previously described (Gilglioni et al., 2018). Lactate is stable in 3% MPA at 4°C and was assayed directly in the MPA extracts.

**Determination of ATP, ADP, and AMP by high performance liquid chromatography (HPLC) for the cytosolic adenylate energy charge**

HepG2 cells were treated with 0.1% DMSO or 50 µM sAC-specific inhibitor LRE1 in HBSS containing indicated substrates. After indicated period of incubation, cytosolic metabolites were extracted with permeabilization buffer (120 mM KCl, 10 mM NaCl, 5 mM EDTA, 20 mM HEPES, pH 7.1) containing 100 µg/mL digitonin. Two hundred microliter of digitonin extracts were mixed with 16 µL 70% perchloric acid for deproteinization and subsequently neutralized with 2.5 M $K_2CO_3$. The neutralized perchloric acid extracts of HepG2 cells were then analysed for AMP, ADP and ATP by high-performance liquid chromatography using a Partisphere SAX column (Whatman International Ltd.) exactly as described (Bontekoe et al., 2017). Adenylate energy charge was defined as ([ATP] + 0.5 × [ADP]) / ([ATP] + [ADP] + [AMP]) according to Atkinson and Walton (Atkinson and Walton, 1967).

**Sodium dodecyl sulfate–polyacrylamide gel electrophoresis (SDS-PAGE) and Western blotting**

Protein concentrations of whole cell lysates in RIPA buffer were quantified by bicinchoninic acid (BCA) assay. Equal amount of protein (40-50 µg) was subjected to SDS-PAGE, transferred to polyvinylidene difluoride (PVDF) membranes by semi-dry blotting and blocked overnight in 5% non-fat milk / PBST (phosphate-buffered saline with 0.05% (w/v) Tween 20) at 4°C. For immunodetection, the PVDF membranes were incubated with primary antibody for 1 hour, washed 3 times with TBST (Tris-buffered saline with 0.05% (w/v) Tween 20), incubated with horseradish peroxidase-conjugated secondary antibody for 1 hour, and washed again 4 times with TBST. All antibodies were diluted in 1% non-fat milk-TBST and incubation was performed at room temperature. The PVDF membrane was developed with homemade enhanced chemiluminescence reagents (100 mM Tris-HCl pH 8.5, 1.25 mM luminol, 0.2 mM p-coumarin and freshly added 3 mM $H_2O_2$) and detected using the ImageQuant LAS 4000 (GE Healthcare Life Sciences). Please refer to Table S3 for the list of primary and secondary antibodies and dilution.

**Statistics**

All results are given as mean ± standard deviation (SD). Statistical significance was determined by two-tailed Student's t-test, one-way analysis of variance (ANOVA) with Turkey's or Dunnett's multiple comparison test, or two-way ANOVA with Sidak's multiple comparison test as indicated in the legends. Statistical analysis was performed with GraphPad Prism 7 (GraphPad Software, La Jolla, CA) with an α error of 0.05.

### Supplementary Figure Legends

**Figure S1. Soluble adenylyl cyclase is an acute switch for aerobic glycolysis that maintains energy homeostasis (related to Figure 1).**

HepG2 cells were preincubated for 1 hour in HBSS for ambient air in the presence of 125 µM octanoate and subsequently transferred to the Seahorse Flux Analyzer. ATP production rates by oxidative phosphorylation ($J_{ATP\_OxPhos}$, A), tricarboxylic acid cycle ($J_{ATP\_TCA}$, B), mitochondria ($J_{ATP\_Mitochondria}$ , C), glycolysis ($J_{ATP\_Glycolysis}$, D), and total ATP production rate ($J_{ATP\_Total}$, E) were derived from OCR and ECAR measurements shown in Figure 1C and 1D as described in *Methods*. (F) HepG2 cells were treated with 0.1% DMSO (vehicle control) or 50 µM LRE1 for 10 and 90 minutes. AMP, ADP, and ATP were measured for the calculation of the adenylate energy charge. Data represent mean ± SD of triplicate determination.

**Figure S2. sAC- and tmAC-derived cAMP have opposite effects on glycogen homeostasis (related to Figure 2)**

HepG2 cells were acutely incubated in glucose-free HBSS with 0.1% DMSO (n = 3), 50 µM LRE1 (sAC-specific inhibitor, n = 3), and 1 µM forskolin (n = 3) for 60 min. The lactate concentration in the medium was determined in samples taken after 15, 30 and 60 minutes of incubation. Data represent mean ± SD of triplicate samples. Results shown are representative of 2 independent experiments.

**Figure S3. Inhibition of sAC-Epac1 signaling induces glycogenolysis in various cell lines (related to Figure 3)**

HeLa cells (A), Caco-2 cells (B), HEK293T cells (C) and Madin-Darby canine kidney (MDCK) cells (D) were pre-incubated for 15 min in glucose-free DMEM and then the medium was refreshed with glucose-free DMEM containing 0.1% DMSO (vehicle control), 50 µM LRE1, or 50 µM (R)-CE3F4. Cells were incubated for 45 minutes and then the supernatant was sampled for lactate determination and cell lysates were prepared for glycogen determination. The cellular glycogen content was expressed as % of the baseline value: 349 ± 6 (n = 4) for HeLa cells (A), 298 ± 23 (n = 4) for Caco-2 cells (B), 34 ± 14 (n = 4) for HEK293T cells (C), 36 ± 0.4 (n = 4) for MDCK cells (D). Statistical analysis: One-way ANOVA with Dunnett's multiple comparisons test (against DMSO group). n.s., not significant; *P<0.05, **P<0.01, ***P<0.001.

**Figure S4. Various immunoblots in HepG2 cells and H69 cholangiocytes (related to Figure 4)**

(A) HepG2 cells were treated as described in the legend to Figure 4D. Activity of AMPK was measured by determining the phosphorylation of its substrate acetyl-CoA carboxylase (ACC) at residue Ser-80 by immunoblotting. (B) H69 cholangiocytes were treated as described for HepG2 cells in the legend to Figure 4D. Phosphorylation of Ser-15 of liver form glycogen phosphorylase (PYGL) was examined by immunoblotting. (C) H69 cholangiocytes were treated as described for HepG2 cells in the legend to Figure 4E. Phosphorylation of Ser-15 of PYGL was examined by immunoblotting.

### Reference for Supplementary materials


Atkinson, D.E., and Walton, G.M. (1967). Adenosine triphosphate conservation in metabolic regulation. Rat liver citrate cleavage enzyme. J Biol Chem *242*, 3239-3241.

Bontekoe, I.J., van der Meer, P.F., van den Hurk, K., Verhoeven, A.J., and de Korte, D. (2017). Platelet storage performance is consistent by donor: a pilot study comparing "good" and "poor" storing platelets. Transfusion *57*, 2373-2380.

Chang, J.C., Go, S., de Waart, D.R., Munoz-Garrido, P., Beuers, U., Paulusma, C.C., and Oude Elferink, R. (2016). Soluble Adenylyl Cyclase Regulates Bile Salt-Induced Apoptosis in Human Cholangiocytes. Hepatology *64*, 522-534.

Chang, J.C., Go, S., Gilglioni, E.H., Duijst, S., Panneman, D.M., Rodenburg, R.J., Li, H.L., Huang, H.L., Levin, L.R., Buck, J., et al. (2021). Soluble adenylyl cyclase regulates the cytosolic NADH/NAD(+) redox state and the bioenergetic switch between glycolysis and oxidative phosphorylation. Biochim Biophys Acta Bioenerg *1862*, 148367.

Gilglioni, E.H., Chang, J.-C., Duijst, S., Go, S., Adam, A.A.A., Hoekstra, R., Verhoeven, A.J., Ishii-Iwamoto, E.L., and Oude Elferink, R.P.J. (2018). Improved oxygenation dramatically alters metabolism and gene expression in cultured primary mouse hepatocytes. Hepatology Communications *2*, 299-312.

Grubman, S.A., Perrone, R.D., Lee, D.W., Murray, S.L., Rogers, L.C., Wolkoff, L.I., Mulberg, A.E., Cherington, V., and Jefferson, D.M. (1994). Regulation of intracellular pH by immortalized human intrahepatic biliary epithelial cell lines. Am J Physiol *266*, G1060-1070.

Mookerjee, S.A., Gerencser, A.A., Nicholls, D.G., and Brand, M.D. (2017). Quantifying intracellular rates of glycolytic and oxidative ATP production and consumption using extracellular flux measurements. J Biol Chem *292*, 7189-7207.

Mookerjee, S.A., Goncalves, R.L., Gerencser, A.A., Nicholls, D.G., and Brand, M.D. (2015). The contributions of respiration and glycolysis to extracellular acid production. Biochim Biophys Acta *1847*, 171-181.
