## Supplementary Tables for "Opposite regulation of glycogen metabolism by cAMP produced in the cytosol and at the plasma membrane"

**Table S1. Formulation of the modified Hank's balanced salt solutions (HBSS) for ambient air.**

| Component | Molecular Weight | Final Conc. (mg/L) | Final Conc. (mM) |
|---|---|---|---|
| $CaCl_2$•$2H_2O$ | 147.01 | 186 | 1.27 |
| KCl | 74.55 | 340 | 4.56 |
| $NaH_2PO_4$•2H2O | 156.02 | 70 | 0.45 |
| $MgSO_4$•$7H_2O$ | 246.47 | 100 | 0.41 |
| $MgCl_2$•$6H_2O$ | 203.30 | 80 | 0.39 |
| NaCl | 58.44 | 7550 | 129.19 |
| $NaHCO_3$ | 84.01 | 0 | 0.00 |
| $Na_2HPO_4$•$2H_2O$ | 177.99 | 60 | 0.34 |
| HEPES-NaOH, pH7.4 | 238.30 | 4766 | 20.00 |

**Table S2. Parameters used for the estimation of ATP production rate in cells that are deprived of extracellular glucose and fueled solely with octanoate. Endogenous glycogen serves as the primary substrate for glycolysis.**

| | | | *max* P/O | | | |
|---|---|---|---|---|---|---|
| **Substrate** | **ATP/Lactate** | ***max*** **$H^+/O_2$** | **Glycolysis** | **TCA, β-ox** | **Oxphos** | **Total** |
| Octanoate | 1.45 | 0.64 | 0 | 0.121 | 2.38 | 2.500 |

**Table 3. List of primary and secondary antibodies used in immunoblotting.**

| Antibodies | Company | Catalog No. | Host | Isotype | Dilution |
|---|---|---|---|---|---|
| ***Primary antibodies*** | | | | | |
| Anti-ACC | Cell signaling | 3676 | Rabbit mAb | IgG | 1:1000 |
| Anti-pACC (Ser80) | Cell signaling | 3361 | Rabbit pAb | IgG | 1:1000 |
| Anti-PYGL | NovusBio | NBP2-32246 | Rabbit pAb | IgG | 1:1000 |
| Anti-phospho-PYGL (Ser15) | MRC-PPU, University of Dundee | S961A | Sheep pAb | NA | Used at 1 µg/mL |
| ***Secondary antibodies*** | | | | | |
| Anti-rabbit IgG, HRP-conjugated | Bio-Rad | 170-6515 | Goat pAb | Anti-serum | 1:3000 |
| Anti-sheep IgG, HRP-conjugated | Sigma | SAB370071 | Rabbit pAb | IgG | 1:1000 |
